# Rewiring the specificity of extra-cytoplasmic function sigma factors

**DOI:** 10.1101/2020.06.23.168245

**Authors:** Horia Todor, Hendrik Osadnik, Elizabeth A. Campbell, Kevin S. Myers, Timothy J. Donohue, Carol A. Gross

## Abstract

Bacterial genomes are being sequenced at an exponentially increasing rate, but our inability to decipher their transcriptional wiring limits our ability to derive new biology from these sequences. *De novo* determination of regulatory interactions requires accurate prediction of regulators’ DNA binding and precise determination of biologically significant binding sites. Here, we address these challenges by solving the DNA-specificity code of extra-cytoplasmic function sigma factors (ECF σs), a major family of bacterial regulators, and determining their regulons. We generated an aligned collection of ECF σs and their promoters by leveraging the auto-regulatory nature of ECF σs as a means of promoter discovery and analyzed it to identify and characterize the conserved amino acid – nucleotide interactions that determine promoter specificity. This enabled *de novo* prediction of ECF σ specificity, which we combined with a statistically rigorous phylogenetic foot-printing pipeline based on precomputed orthologs to predict the direct targets of ∼67% of ECF σs. This global survey indicated that ECF σs play varied roles: some are global regulators controlling many genes throughout the genome that are important under many conditions, while others are local regulators, controlling few closely linked genes in response to specific stimuli. This analysis reveals important organizing principles of bacterial gene regulation and presents a conceptual and computational framework for deciphering gene regulatory networks.

## INTRODUCTION

The genomes of medically, industrially, and environmentally important bacteria are being sequenced at a rapidly increasing rate, and automatic annotation of genes has become relatively straight-forward. However, the utility of these sequences is limited by our inability to decipher their transcriptional wiring. Because adaptation to new environments or conditions is driven both by changes in the regulation of existing genes and by acquisition of novel functions, deciphering gene regulation from genome sequences is a key aspect of understanding cellular lifestyles, the evolution of pathogenesis, intrinsic antibiotic resistance, and biofilm growth.

In bacteria, a major point of transcriptional regulation is the use of sigma factors (σs) to direct RNA polymerase (RNAP) to specific promoters (Gruber and Gross 2003). A vast majority of σs belong to the σ^70^ family, which consists of four phylogenetically and structurally related groups (Paget 2015). The Group 1 “housekeeping” σs are highly conserved, universally essential, and responsible for recognizing 1000s of promoters in each bacterial genome. The Group 2-4 alternative σs are active under specific growth or environmental conditions and effect specialized transcriptional programs by directing RNAP to a smaller set of distinct promoters. Group 4 σs, also known as <u>e</u>xtra-<u>c</u>ytoplasmic function (ECF) σs, are found in all major bacterial phyla and are the most abundant and diverse group of alternative σs, representing 5-10% of an average genome’s regulatory repertoire (Staroń et al. 2009). They have been implicated in the regulation of genes involved in differentiation, metal homeostasis, outer membrane integrity, oxygen response, and many other processes (Kelemen et al. 1996; Braun, Mahren, and Ogierman 2003; Mecsas et al. 1993; Donohue 2019). Despite their importance, the vast majority of ECF σs remain uncharacterized and no computational method exists for determining the set of genes regulated by an ECF σ (its regulon).

ECFσs are small proteins (∼200 amino acids) with two well conserved globular domains, σ4 and σ2, that interact respectively with the -35 and -10 regions of the core promoter (Gruber and Gross 2003). Despite their small size and divergent sequence, ECF σs fulfill the functional and structural requirements of all σs: (1) recruiting the RNA-polymerase core enzyme, (2) recognizing the promoter, and (3) melting the dsDNA to nucleate the transcription bubble. However, because they lack a σ3 domain and because their diverged σ2 domain lacks the aromatic residues important for promoter melting (Koo et al. 2009; Campagne et al. 2014), ECF σs are less proficient at promoter melting than other σs, resulting in a requirement for near consensus promoters. Additionally, many ECF σs auto-regulate their own promoter, amplifying activating cues and enabling a strong transcriptional response despite their low promoter melting ability (Staroń et al. 2009). The small size, requirement for a near-consensus promoter, and auto-regulatory nature of many ECF σs make them amenable to computational approaches that may be able to provide insights into the promoter recognition properties of these σs.

The auto-regulatory nature of many ECF σs has been used as a means of promoter discovery (Staroń et al. 2009; Rhodius et al. 2013). Assuming that similar protein sequence implies similar promoter motifs, Mascher and colleagues (Staroń et al. 2009) separated ECF σs into 43 groups and identified the putative autoregulatory promoter of 18 of them by examining their upstream regions for over-represented bi-partite motifs. By expanding this search to regions upstream of ECF σ containing operons and divergently transcribed genes, Rhodius and colleagues (Rhodius et al. 2013) identified the putative promoter motifs of an additional 11 ECF σ groups. However, the ECF σ promoter motifs they discovered were not suitable for determining individual ECF σ regulons. First, many of the 43 ECF σ groups were quite broad and contained multiple ECF σs from the same genome. Since it is unlikely that multiple ECF σs in a single bacterium regulate identical regulons, these promoter motifs are likely an ensemble of multiple distinct promoters. Second, promoter motifs can be found only for auto-regulated ECF σs, precluding the determination of non-auto-regulatory ECF σ regulons. Finally, even with accurate knowledge of the -35 and -10 motifs, identifying the true regulon is complicated by the large fraction of false positive sites (Urtecho et al. 2020; Huerta and Collado-Vides 2003; Rhodius, Mutalik, and Gross 2012).

We overcame these hurdles by computationally and experimentally analyzing the protein determinants of promoter recognition to elucidate how amino acid identities at key positions determine the promoter specificity of ECF σs. This ECF σ-specific code enables the universal prediction and rational re-engineering of ECF σ promoter specificity. To take advantage of our ability to individually predict the promoter specificity of all ECF σs, we developed a statistically rigorous phylogenetic foot-printing pipeline based on eggNOG orthology groups (Huerta-Cepas et al. 2016) to discover ECF σ regulons. Using these tools, we successfully predicted the regulons of ∼67% of bacterial ECF σs. We found that ECF σs regulate a broad range of functions and can be generally classified into “global” and “local” ECF σs. Whereas “local” ECF σs have small regulons and a sparse distribution within clades, “global” ECF σs regulate many promoters and are found in most members of a bacterial clade. Our analysis reveals important organizing principles of bacterial regulatory network evolution and provides the first truly global survey of gene regulatory interactions in bacteria.

## RESULTS AND DISCUSSION

### ECF σs interact with DNA in a conserved fashion

Predicting and rationally engineering the specificity of a DNA binding protein requires conserved interactions between specific positions in the protein and the DNA. To determine whether ECF σs meet this requirement, we constructed a large database of ECF σs, determined their putative auto-regulatory promoters, and performed a mutual information (MI) analysis to determine whether specific amino acid positions co-varied with specific nucleotide positions across evolutionary time.

We first identified ∼133,000 ECF σs in ∼20,000 genomes (Supplementary Table 1) from the proGenomes database (Mende et al. 2017) using Pfam models for the σ2 and σ4 domains, as previously described (Staroń et al. 2009). This initial set was reduced to 84,009 ECF σs in 10,000 genomes by eliminating ECF σs with identical upstream and protein sequences. To determine the putative auto-regulatory promoter motifs of these ECF σs, we clustered them based on their protein sequences (Methods) and used BioProspector (X. Liu, Brutlag, and Liu 2001) to search for overrepresented motifs in the upstream sequences of each ECF σ cluster (Figure 1A). We maximized our chances of identifying motifs by applying two distinct clustering methods (K-mer distance and eggNOG identifier; Supplementary Table 1; Huerta-Cepas et al. 2016), searching both 150bp and 300bp upstream of the ECF σ translation start site, and considering sequences upstream of the ECF σs, upstream of the operons encompassing the ECF σs (if applicable), and upstream of the divergently transcribed gene (if applicable), similar to previous work (Rhodius et al. 2013). Using these different parameter sets, we identified 91,552 putative promoters belonging to 41,665 unique ECF σs. Motifs were aligned and each putative promoter sequence was associated with its respective ECF σ sequence, generating a matched collection of aligned promoters and ECF σ sequences (Supplementary Table 2).

**Figure 1.**
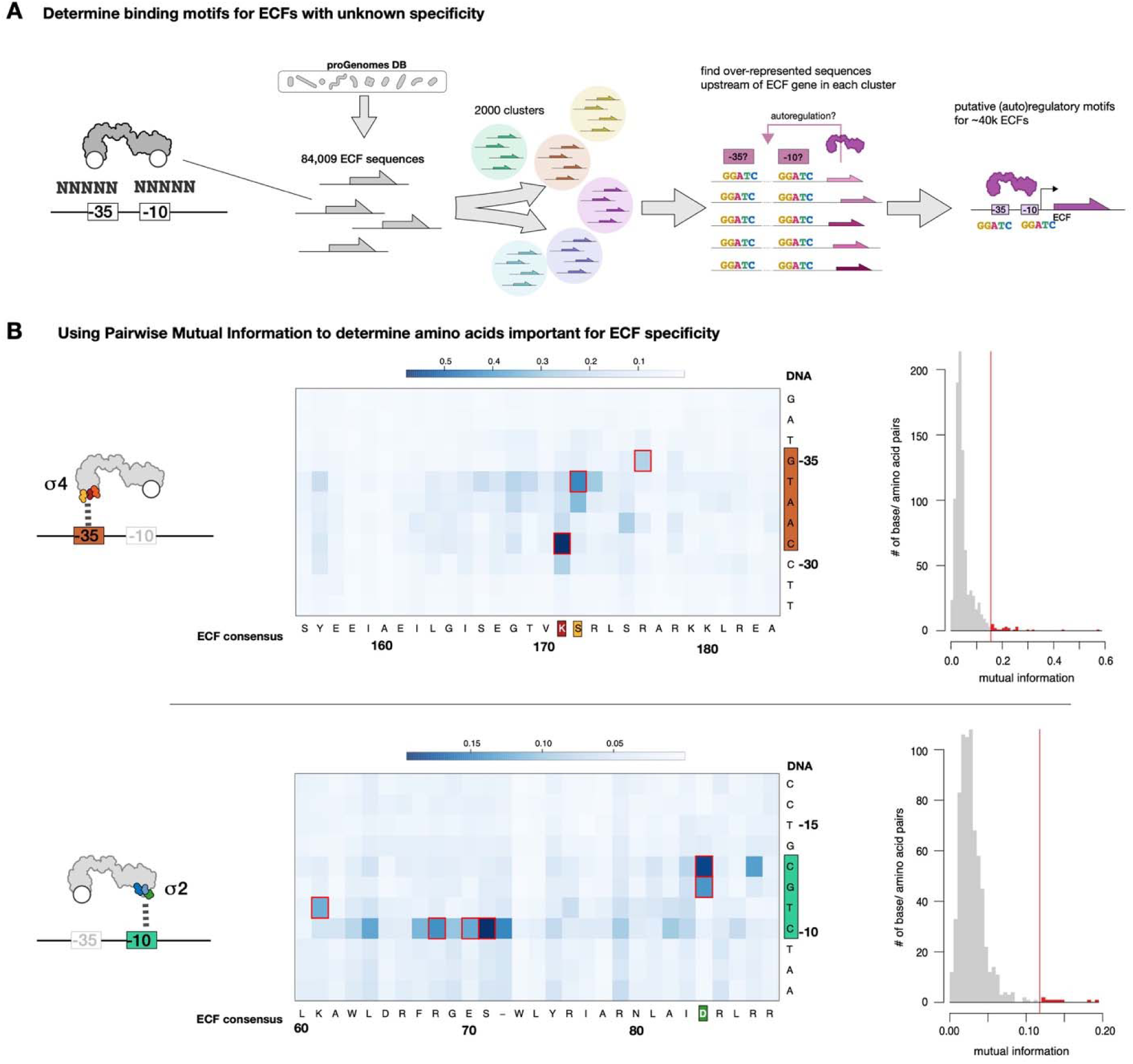
(A) Schematic depicting the process of identifying putative ECF σ motifs (B) Mutual information maps and distributions for the interaction between σ4 domain and -35 & for the interaction between σ2 and -10.

We next performed a MI analysis to determine whether specific amino acid positions co-varied with specific promoter positions. MI is a measure of the co-variation between two variables (in this case, amino acid and nucleotide identity). If a certain amino-acid position interacts with a certain nucleotide position in the promoter in most ECF σs, the identity of the base at that position should co-vary with the amino acid position, resulting in high MI. In contrast, amino-acid positions and nucleotide positions which do not consistently interact would not co-vary, resulting in low MI. We calculated the MI between all positions in the promoter and protein alignments and found that only a few amino acids – nucleotide positions had high MI (Figure 1B). High MI amino acid – nucleotide interactions were consistent with specificity determining interactions observed in the structures of 3 divergent ECF σs bound to their cognate promoters: *E. coli* RpoE and *Mycobacterium tuberculosis* SigH and SigL (Lane and Darst 2006; Campagne et al. 2014; Lin et al. 2019; Fang et al. 2019; Li et al. 2019; Figure 1B). Taken together, these data suggest that the mechanism of ECF σ– promoter interaction is conserved in a majority of ECF σs despite their extensive sequence divergence, implying that our collection of matching ECF σs and putative promoters can serve as a dictionary in which the promoter specificity conferred by any amino acid at key DNA recognizing positions can be determined. For example, to determine the -31 base specificity of a specific ECF σ, we determine the identity of the amino acid in our ECF σ of interest equivalent to Arg171 in *E. coli* RpoE (hereafter, ECF σ positions are referred to by the equivalent position in *E. coli* RpoE, e.g. Arg171^RpoE^). We then assess the corresponding -31 base identity of all natural ECF σs in our database that have the same amino acid identity and ascribe that -31 base preference to the ECF σ of interest. By aggregating the promoter specificity for individual promoter positions conferred by the amino acids at key positions in this manner, the complete promoter specificity of any ECF σ can, in theory, be deduced.

### The identify of key amino acid positions determines promoter specificity

Our MI analysis identified key amino acid positions in the σ2 and σ4 domains of ECF σs that appear to determine the promoter specificity. This analysis implies that single amino acid substitutions at these positions should affect the promoter specificity of an ECF σ in a predictable way, enabling the design of ECF σs with arbitrary promoter specificity as well as prediction of natural ECF σ promoters. To determine whether this is the case, we experimentally characterized the effect of all possible substitutions at 3 key amino acid positions on promoter specificity using a previously described system for ECF σs expression and promoter activity measurement (Rhodius et al., 2013; Figure 2A).

**Figure 2.**
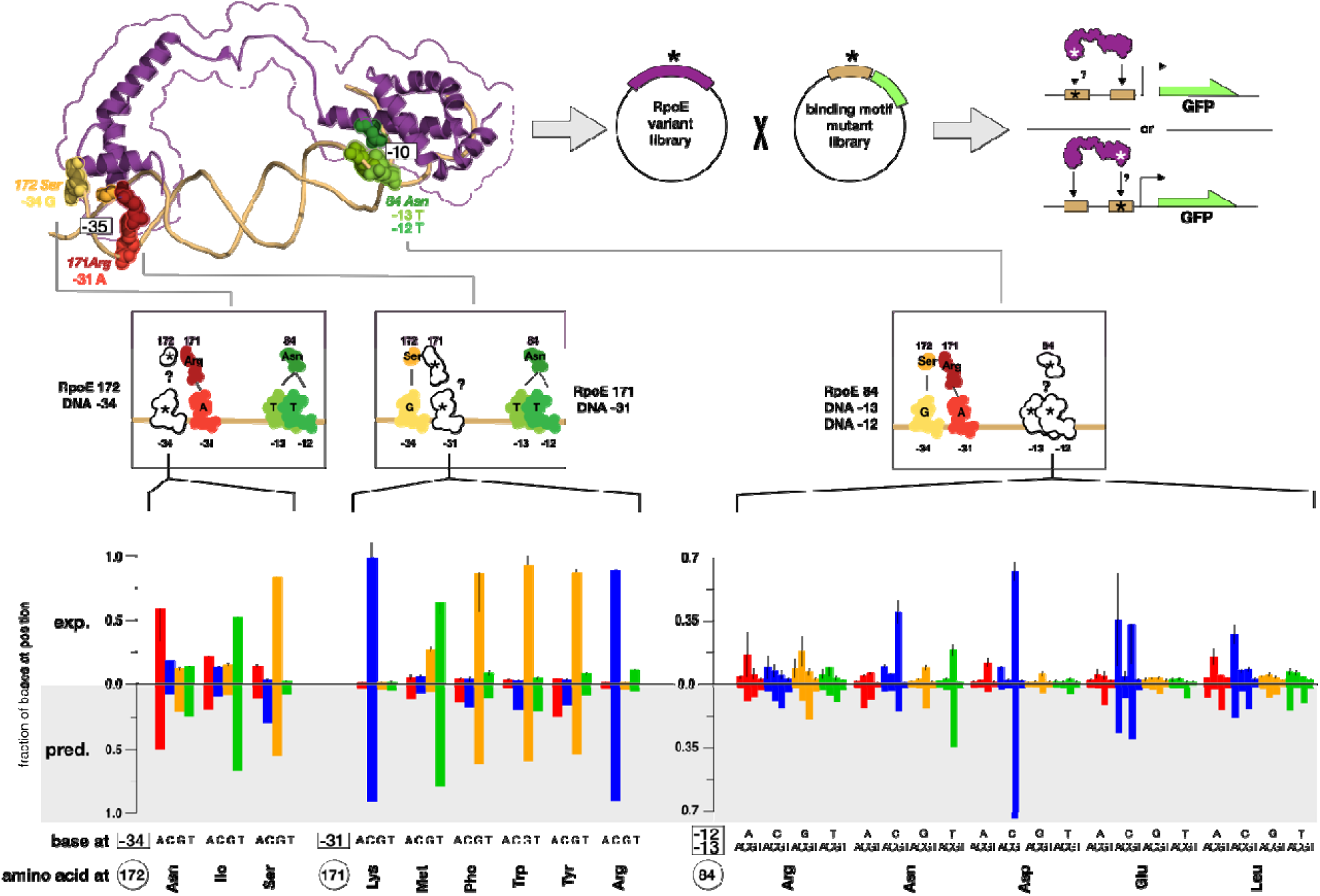
Comparison of experimental and predicted effects of single amino-acid mutations on promoter specificity.

We tested how the identity of the Arg171^RpoE^ amino acid affects -31 nucleotide specificity by generating all natural amino acid substitutions at the Arg171^RpoE^ position of ECF32_1122 and testing the ability of these mutant ECF σs to drive transcription of GFP *in vivo* from pECF32 promoter constructs containing every possible -31 nucleotide (Rhodius et al. 2013); Methods; Supplementary Table 3). Mutations at Arg171^RpoE^ drastically altered -31 specificity (Figure 2). For example, whereas the wild-type ECF32_1122 (containing arginine at the mutagenized position) specifically recognized a C at the -31 position, mutation to methionine changed specificity to T, while mutation to tyrosine changed specificity to G. Importantly, the experimentally determined -31 nucleotide preference of most active ECF σ mutants agreed strongly with the -31 promoter identity of naturally occurring ECF σs with same Arg171^RpoE^ amino acid in our dataset (R^2^ = 0.57; *p <* 10^−10^; 14/20 active).

We similarly tested the effect of substitutions at the Ser172^RpoE^ position on -34 specificity using ECF14_1324 and its cognate promoter pECF14 (Rhodius et al. 2013; Methods; Supplementary Table 3). DNA bound ECF σ structures (Lane and Darst 2006; Fang et al. 2019; Li et al. 2019) and our MI analysis suggest that -34 is recognized through a multipartite interaction driven primarily by the amino acid at the Ser172^RpoE^ position but with contributions from the amino acids at the Gly168^RpoE^ and Thr169^RpoE^ positions. Consistent with multipartite determination of - 34 specificity, we found that only 4/20 Ser172^RpoE^ substitutions were active. Despite this, ECF14_1324 mutants with active substitutions showed substantial changes in -34 specificity (Figure 2) that were well correlated to the specificity of naturally occurring ECF σs with the same Ser172^RpoE^ identity in our dataset (R^2^ = 0.37; *p* < 0.02). Our MI analysis identified Arg173^RpoE^ as a further determinant of -34 specificity. Since the amino acid at Arg173^RpoE^ is not positioned to make base-specific interactions, this is likely due to a role in positioning the amino acid at Ser172^RpoE^. Consistent with this hypothesis, an even stronger correlation between experimentally determined -34 specificity and the -34 specificity of natural ECF σs (R^2^ = 0.47; *p* < 0.005) was observed when both the Ser172^RpoE^ and Arg173^RpoE^ positions were considered. Finally, we tested the effect of amino acid substitutions at the Asn84^RpoE^ position of ECF11_987 on -12 and -13 nucleotide specificity (Supplementary Table 3). Whereas all previously solved DNA-bound ECF σ structures reported that the amino acid at the Asn84^RpoE^ position makes a hydrogen bond with the -12 base of the promoter (Campagne et al. 2014; Lin et al. 2019; Fang et al. 2019; Li et al. 2019), our MI analysis suggested that the Asn84^RpoE^ identity determines both -12 and -13 nucleotide specificity. We therefore tested the ability of all Asn84^RpoE^ substitutions of ECF11_987 to drive transcription from 16 variants of the ECF11_987 promoter modified at the -13 and/or -12 nucleotide positions (Methods). We found that amino acid substitutions at Asn84^RpoE^ drastically altered both -12 and -13 promoter specificity (Figure 2), but that some substitutions appear to recognize only the -12 position, whereas others appear to recognize both -12 and -13 positions. Consistent with the crystal structure, an asparagine at the Asn84^RpoE^ position ECF11_987 activated all promoters with a -12G, whereas the aspartic acid mutant primarily activated promoters with -13C and -12G. The altered promoter specificities of our ECF σ mutants were strongly correlated to the promoter specificity of natural ECF σs with the same amino acids at Asn84^RpoE^ (Figure 2; R^2^ = 0.49; *p <* 10^−16^).

Taken together, these comprehensive mutagenesis studies, performed on 3 distinct ECF σs, demonstrate the stringent yet plastic promoter specificity of ECF σs. All 3 ECF σs tested were exquisitely sensitive to both single nucleotide changes to their cognate promoters and single amino acid changes in both the σ2 and σ4 domains. Importantly, the changes in specificity caused by the amino acid mutations correlated strongly with the promoter preferences of natural ECF σs containing those amino acids. This remarkable correlation suggests that key amino acid identities determine promoter specificity with minimal effect from protein context. This enables the *de novo* prediction of ECF σ promoter specificity.

### Modeling studies confirm the importance of novel specificity determinants

Although most high MI interactions identified amino-acid nucleotide pairs previously implicated in promoter recognition, some high MI interactions highlighted interactions not previously thought to play a role in promoter recognition. To better understand how amino acid changes at these positions may affect promoter specificity, we used modeling methods (similar to Kim, Park, and Park 2016) to explore two non-canonical amino acid – nucleotide interactions.

First, MI analysis uncovered a correlation between the identity of the position equivalent to *E. coli* RpoE Phe175 (hereafter referred to as the Phe175^RpoE^ position) and the -32 nucleotide. Looking across our compendium of natural ECF σs and promoters, we found that many natural ECF σs with an arginine at Phe175^RpoE^ had a (non-template strand) T at the -32 promoter position. We therefore modeled both the amino-acid and nucleotide changes onto the *E. coli* RpoE DNA-bound σ4 structure (Lane and Darst 2006). We found that a phenylalanine to arginine substitution at Phe175^RpoE^ would normally lead to the loss of three non-polar interactions with the -32 template strand T methyl, C6, and ribose C2’. The compensatory -32T mutation places the -32 template strand A N7 acceptor near the arginine at Phe175^RpoE^ and likely compensates for the loss of the non-polar interactions with the formation of a H-bond from the guanidine group (Supplementary Figure 1A). This result strongly suggests that mutating Phe175^RpoE^ to arginine is likely to require -32T to maintain activity, consistent with our predictions.

Second, we found a surprising link between the identity of the Lys56^RpoE^ position and the identity of the -11 nucleotide. Previous structural studies implicated Lys56^RpoE^ primarily in interactions with the -12 nucleotide and suggested that the -11T was predominantly recognized by Ile77^RpoE^, Ala60^RpoE^, and Asn80^RpoE^ (Campagne et al. 2014). Much of the co-variation between the amino acid at Lys56^RpoE^ and the nucleotide at the -11 position was driven by the association of acidic amino acids, such aspartic acid and glutamic acid, with an A at -11 and conversely of basic amino acids, such as lysine and arginine, with a T at -11. We therefore modeled the effects of acidic substitutions at the Lys56^RpoE^ position on a pre-existing structure (Campagne et al., 2014). Basic amino acids at the Lys56^RpoE^ position normally form an ionic interaction with O4 of -11T by donating an electron, explaining their affinity for -11T. By contrast, the -11A nucleotide likely donates 2 electrons to acidic substitutions at the Lys56^RpoE^ position, creating a strong (reversed) ionic interaction that explains the correlation seen in the MI analysis (Supplementary Figure 1B).

Taken together, our modeling results strongly support the hypothesis that the sparse and modular correlations observed in the MI analysis are causal drivers of ECF specificity and highlight the importance of previously unappreciated interactions for the promoter specificity of ECF σs.

### *De novo* prediction of ECF promoter specificity

That only a few amino acid positions in ECF σs make sequence specific DNA contacts suggests that *de novo* prediction of promoter specificity based on primary sequence may be attainable. Such an approach would be suitable for inferring the promoter motifs of non-autoregulatory and recently diverged ECF σs, neither of which cannot be accurately predicted through clustering-based approaches.

To predict the promoter specificity of ECF σs, we first selected the 1-2 amino acids most correlated to the identity of each nucleotide position in the promoter based on the results of our MI analysis and published structures (Supplementary Table 4). To determine the promoter sequence specificity of a given ECF σ, we consider each nucleotide position separately. At each promoter position we evaluate the DNA specificity of natural ECF σs that have the same amino acid(s) at the specificity determining positions as our ECF σ of interest. By concatenating these individual sequence preferences, we determine PWMs representing the -35 and -10 sequence specificity of a given ECF σ (Figure 3A). We determine how efficiently a given ECF σ will activate a given promoter by scoring every possible position in the promoter sequence using the -35 and -10 PWMs and a 16bp, 17bp or 18bp spacer as previously described (Rhodius and Mutalik 2010). We do not assess a spacer penalty, because sequence determinants of ECF σ spacer preference are not known.

**Figure 3.**
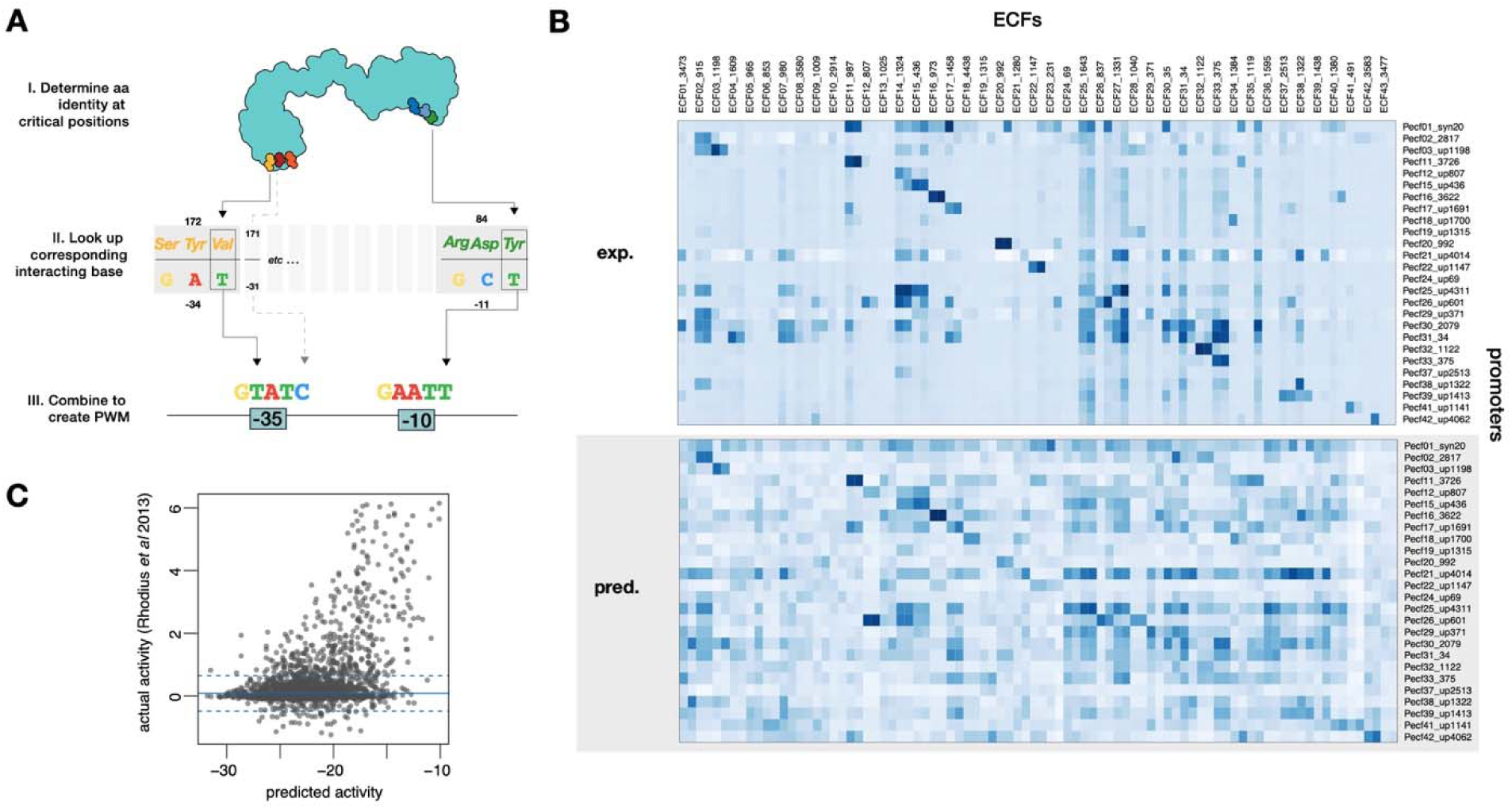
(A) Schematic depicting the prediction of ECF σ motifs based on the identify of amino acids at specific positions. (B) Comparison between the actual (top) and predicted (bottom) activity of 86 ECF σs on 26 promoters, from (Rhodius et al. 2013). Correlation between actual and predicted (C).

To assess the predictive power of this model, we tested our ability to predict 2,236 ECF σs – promoter interactions for which *in vivo* experimental data is available (Rhodius et al. 2013). In that study, two representatives from each of the 43 major ECF σ classes were synthesized and heterogeneously expressed in *E. coli.* Each of these 86 ECF σs was then tested for its ability to drive transcription from 26 promoters constructed from the putative consensus auto-regulatory promoters of the 43 major ECF σ classes (Figure 3B). This rich set of promoter-ECF σ interactions (total of 2,236) contains representatives of many ECF σ classes, diverse promoters, and varying degrees of activation. We predicted the -35 and -10 recognition preferences for each of the 86 ECF σs in this study on the basis of their primary sequence and scored each of the 26 promoter sequences using these PWMs (Figure 3B). Predicted activity of these 86 ECF σs was strongly correlated (R = 0.44; *p* < 10^−16^; Figure 3C; Supplementary Table 5) to their experimentally measured activity across all promoter constructs. Remarkably, this correlation was comparable to that observed when scoring the experimentally determined PWM of *E. coli* RpoE on its native promoters (Rhodius and Mutalik 2010; *in vitro* R = 0.45; *in vivo* R = 0.60). Taken together, these data suggest that the promoter specificity of diverse ECF σs can be accurately predicted from their primary sequence.

### Phylogenetic foot-printing of ECF σ regulons

Having established an accurate *de novo* method for predicting the -35 and -10 promoter specificity of ECF σs, we next sought to determine what genes were regulated by the 84,009 non-duplicate ECF σs in our dataset. Despite accurate predictions of -35 and -10 specificity, previous studies on σ^70^ and *E. coli* RpoE have highlighted the difficulty of predicting functional promoters even when the -35 and -10 PWMs have been empirically determined. For example, scanning the *E. coli* genome using optimized σ^70^ -35 and -10 PWMs recovered 86% of known promoters, but exhibited a false positive rate (FPR) of ∼80% (Huerta and Collado-Vides 2003). Similarly, scanning the *E. coli* genome using experimentally determined RpoE -35 and -10 PWMs and other sequence characteristics correctly identified 88% of RpoE-regulated promoters but had a FPR of 92% (Rhodius et al. 2006). In the context of predicting ECF σ regulons, such a high FPR would obfuscate the true regulons and preclude biological interpretation of their function.

To overcome these high FPR rates and precisely determine which genes may be regulated by a particular ECF σ, we developed a phylogenetic foot-printing approach (Figure 4A). Phylogenetic foot-printing determines true ECF σ binding sites by comparing orthologous DNA sequences from closely related species. Because natural selection maintains functional elements, such as *bona fide* ECF σ binding sites, during evolution, this method can identify genes regulated by an ECF σ of interest with a much lower FPR than other methods. In spite of its advantages, phylogenetic foot-printing is computationally complex and difficult to parametrize. By developing a pipeline based on the precomputed orthologous groups (OGs) from eggNOG 4.5 (Huerta-Cepas et al. 2016) as implemented in the user-friendly proGenomes database (Mende et al. 2017) and using randomized PWMs to determine significance, we overcome these issues and developed a rapid, high-throughput, and statistically sound method of phylogenetic foot-printing (Methods; Figure 4A).

**Figure 4.**
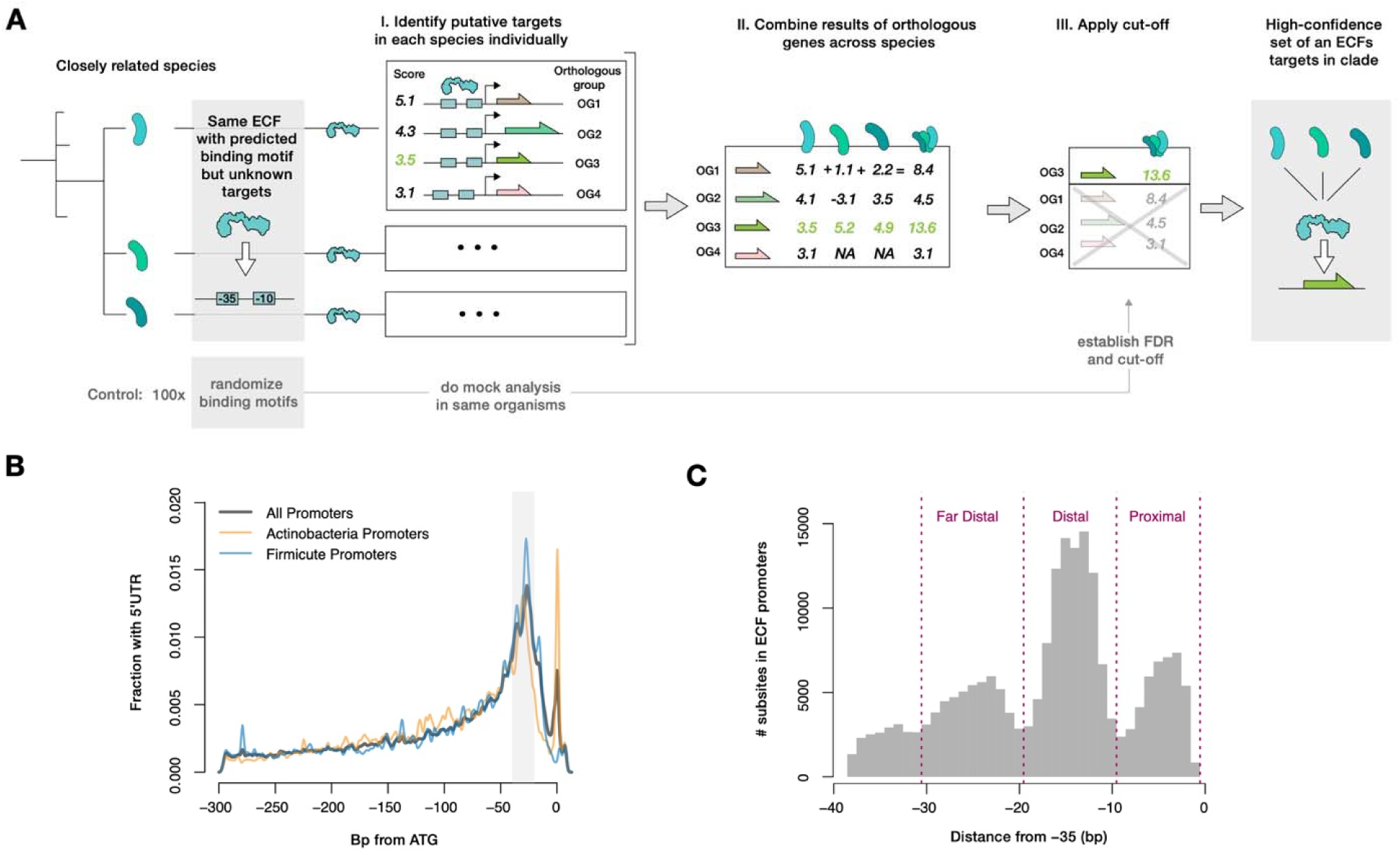
Distribution of 5’UTR lengths in predicted ECF σ promoters in all clades (black), Actinobacteria (orange), and Firmicutes (blue).

We applied this method to all 84,009 non-duplicate ECF σs in our data set and found at least one significantly (FDR < 0.05) regulated eggNOG (effectively, at least one gene) for 67% of the ECF σs. This data-set consists of >600,000 ECF σ-gene interactions occurring at >240,000 specific promoters, and represents the first truly global survey of ECF σ function.

### Predicted ECF σ promoters are similar to experimentally characterized promoters

To assess the accuracy of our predictions, we explored whether these putative ECF σ promoters shared common properties of bacterial promoters. We reasoned that since our phylogenetic foot-printing approach considers only the predicted -35 and -10 specificities, the presence of additional promoter characteristics such as 5’UTR length, initiating nucleotide, and the presence of UP-elements would only arise if biologically relevant and active promoters were being preferentially identified.

We first explored the 5’UTR length of our predicted ECF σ promoters. Previous studies of transcription start sites (reviewed in Beck and Moll 2018) have identified a conserved distribution (Figure 4B) of 5’UTR lengths across bacterial phyla. The distribution of 5’UTR lengths of our predicted ECF σ promoters was similar to this distribution (Figure 4B), exhibiting a peak around ∼20-40bp, which approximately corresponds to the space required for ribosome assembly on the Shine-Dalgarno sequence (Mendoza-Vargas et al. 2009). The distribution of predicted 5’UTR lengths was similar in all bacterial phyla, with the exception of extremely short 5’UTR corresponding to leaderless mRNAs, whose abundance is known to vary within and across bacterial phyla (Figure 4B; Table 1; Beck and Moll 2018).

**Table 1.**
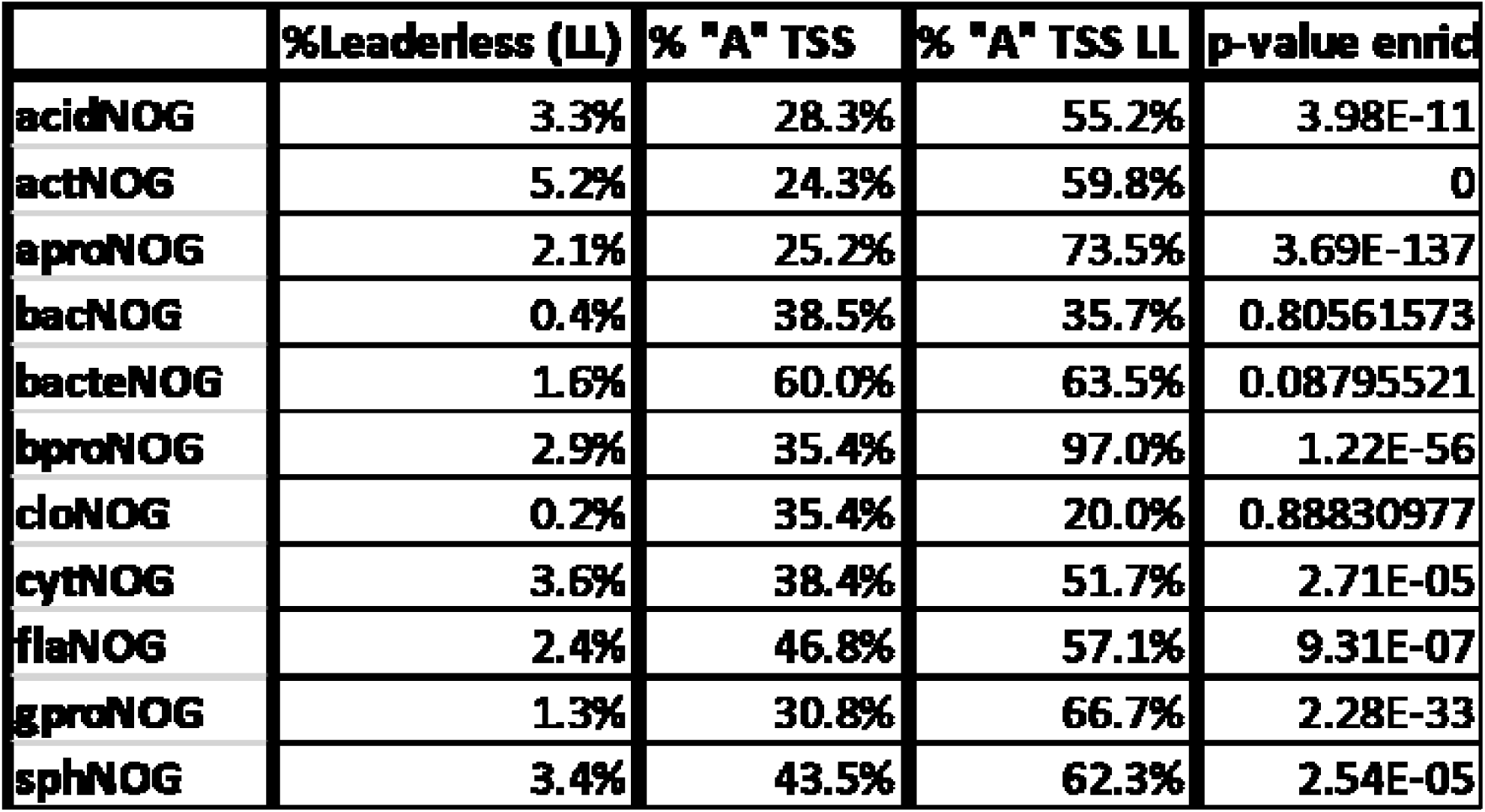
Leaderless transcripts are enriched for “A” TSS.

We next explored the region downstream of the -10 element in predicted ECF σ promoters. Many bacterial promoters contain a pyrimidine at the -1 position of the non-template strand and a purine at the +1 position. This combination allows favorable base-stacking interactions between the -1 template strand purine and the +1 non-template strand purine (Basu et al. 2014). Consistent with this, we found that in 67.3% of putative ECF σ promoters, a pyrimidine-purine sequence was present at one of the two likely transcription start sites (equivalent to the - 1 or +1 position, as counted from the flipped out -10 base). Pyrimidine-purine pairs were enriched relative to their purine-pyrimidine counterparts at the two likely transcription start site positions, but not before or after. Since pyrimidine-purine pairs were present in the approximate TSS for 67.3% of promoters, we assumed that the purine base was the initiating nucleotide (+1) for these promoters. Although both adenine and guanine are commonly used as initiating nucleotides in all phyla, predicted leaderless mRNA promoters were highly enriched for adenine initiating nucleotides (Table 1), as would be expected considering the importance of the AUG codon for recruiting the ribosome to leaderless mRNAs (Beck and Moll 2018).

Finally, we searched for UP-element subsites in predicted ECF σ promoters. UP-elements consist of A/T rich tracts (Estrem et al. 1998) with extremely narrow minor grooves upstream of the -35 element that make sequence specific interactions with the C-terminal domain of one or both alpha subunits of RNAP (αCTD). UP-element subsites must be located on the same face of the DNA-helix as RNAP, but can be located immediately upstream of the -35 element (proximal UP-element subsite), one DNA-helix turn upstream (distal UP-element subsite) or two DNA-helix turns upstream (far distal UP-element subsite). To determine whether our predicted ECF σ promoters had UP-elements, we predicted DNA shape upstream of all ∼240,000 predicted ECF σ promoters and defined putative UP-element subsites as two adjacent nucleotides with a minor groove width >3.5A. We found that ∼25% of putative ECF σ promoters contained at least one UP-element subsite. UP-element subsites were preferentially located at the proximal, distal, and far-distal subsites (Figure 4C), and the frequency of UP-elements varied by bacterial phyla, consistent with previous reports (Table 2; Hubin et al. 2017).

**Table 2.** Distal UP-elements common in ECF promoters.

|  | **Total %** | **% 16bp** | **% 17bp** | **% 18bp** | **% Proximal** | **% Distal** | **% Far Distal** |
| --- | --- | --- | --- | --- | --- | --- | --- |
| **acidNOG** | 9.0% | 12.2% | 7.6% | 6.6% | 2.7% | 4.1% | 2.0% |
| **actNOG** | 3.2% | 1.8% | 3.8% | 2.2% | 0.5% | 2.0% | 0.8% |
| **aproNOG** | 11.3% | 10.3% | 13.7% | 7.2% | 1.8% | 6.9% | 2.9% |
| **bacNOG** | 37.3% | 38.4% | 40.0% | 26.2% | 13.5% | 16.5% | 12.5% |
| **bacteNOG** | 49.6% | 47.2% | 60.6% | 30.1% | 6.4% | 36.7% | 18.1% |
| **bproNOG** | 22.0% | 21.6% | 24.6% | 13.4% | 4.7% | 16.4% | 3.7% |
| **cloNOG** | 43.9% | 40.4% | 50.0% | 36.5% | 14.9% | 24.1% | 16.7% |
| **cytNOG** | 34.0% | 36.0% | 36.0% | 26.9% | 7.6% | 19.6% | 11.4% |
| **flaNOG** | 49.6% | 53.4% | 51.6% | 32.4% | 14.9% | 27.4% | 18.2% |
| **gproNOG** | 14.9% | 16.3% | 16.0% | 9.5% | 4.8% | 8.0% | 3.5% |
| **sphNOG** | 40.2% | 38.8% | 46.1% | 28.0% | 8.5% | 22.6% | 14.1% |
| **Overall** | **24.7%** | **30.9%** | **23.6%** | **15.6%** | **6.9%** | **13.6%** | **8.3%** |

These data demonstrate that our predicted ECF σ promoters share many characteristics with experimentally determined promoters, including 5’UTR length, pyrimidine-purine pairs near the start site, and UP-elements upstream of the -35. Taken together these data suggest that our phylogenetic foot-printing approach correctly identifies biologically relevant and active bacterial promoters. Moreover, the strength of these signals indicates that our pipeline achieves a low false positive rate when identifying promoters, a prerequisite for biological interpretation.

### ECF σ regulons are accurately predicted

Given that our set of putative ECF σ promoters share many properties with active bacterial promoters, we next assessed whether the specific genes predicted to be regulated were correct. Although no regulon information exists for an overwhelming majority of ECF σs in our dataset, approximately 50% of ECF σs have been reported to auto-regulate (Staroń et al. 2009; Rhodius et al. 2013; Figure 1). Taking advantage of this positive control, we therefore assessed what fraction of ECF σs are predicted to autoregulate by our pipeline. We found that 52.9% of ECF σs with significant regulons (*p* < 0.05) included themselves in their regulon, a percentage which did not vary substantially when the *p-*value threshold for significance was raised (55.3% autoregulatory at *p* < 0.01). Among ECF σs for which we identified auto-regulatory promoters, As expected, ECF σs for which auto-regulatory promoters were previously identified (Figure 1) were more likely to auto-regulate than ECF σs for which we did not previously identify auto-regulatory promoters (70% for ECF σs with previously identified auto-regulatory promoters, 33% for other ECF σs), highlighting the accurate and precise regulons determined by combining our DNA-specificity predictions and phylogenetic foot-printing pipeline.

To more specifically probe the accuracy of regulon predictions, we turned to publicly available gene expression data for 13 diverse bacterial species. We reasoned that the expression of genes in true ECF σ regulon, likely including the ECF σ itself, should be correlated across experimental conditions. Therefore, we assessed the accuracy of our regulon predictions for each ECF σ in these 13 species by comparing the correlation between the expression of the predicted ECF σ regulon and the same number of randomly selected genes across all publicly available gene expression data from NCBI GEO (Myers et al. 2020). We found that for 41% of the predicted ECF σ regulons examined (37 out of 90; Supplementary Table 6), the gene expression profiles were significantly more correlated than expected by random chance, suggesting that these predicted regulons represent co-regulated genes. The lack of significant correlations for the remaining ECF σs may be due to incorrect regulon predictions, an insufficiently diverse set of gene expression data (i.e. no conditions that activated the σ), or small but correctly predicted regulons that fail to reach the threshold of statistical significance. Consistent with the idea that small regulons may be more difficult to validate using this method, statistically significant predicted regulons were significantly larger (15.7 genes) than non-significant regulons (7.3 genes; *p <* 0.01). Taken together these data strongly suggest that our model of ECF σ promoter specificity combined with our phylogenetic foot-printing approach to regulon determination correctly identifies the promoter regions and members of ECF σ regulons.

### ECF σs function as both global and local regulators

Studies of genome-wide gene regulatory networks in eukaryotes (Luscombe et al. 2004), archaea (Bonneau et al. 2007), and bacteria (Martínez-Antonio and Collado-Vides 2003) have identified hierarchical network structures consisting of global and local transcriptional regulators. These differ in the number of regulated genes, their conservation, and the diversity of conditions in which they exert their effects. Most well characterized ECF σs such as RpoE in the γ-proteobacteria, SigR/SigH in the Actinobacteria, and EcfG in α-proteobacteria are global regulators that control the expression of many genes in response to diverse conditions (Rhodius et al. 2006; Park, Lee, and Roe 2019; Fiebig et al. 2015). To determine whether a majority of ECF σs function as global regulators, we first quantified the number of promoters regulated by each ECF σ with significant predictions in our dataset. Known global regulators had large average predicted regulons (regulated promoters: RpoE - 10, SigR/SigH - 31, EcfG - 19), however, the number of promoters regulated by ECF σs varied greatly, from 1 to 85 promoters (Figure 5). Nearly half of ECF σs in our dataset (47.8%) were predicted to regulate 3 or fewer promoters. We eliminated the two most obvious technical explanations for this observation, small ECF σs clusters or low information motif predictions, by assessing the correlation between these factors and predicted regulon size. Although both motif information content and cluster size were significantly correlated to regulon size (all *p* < 10^−16^), together these parameters only explained a small amount of the variability in regulon size (R^2^ < 0.1), suggesting that much of the variation in regulon size is biologically meaningful. We therefore sought to determine if the ECF σs predicted to have large regulons are *bona fide* global regulators.

**Figure 5.**
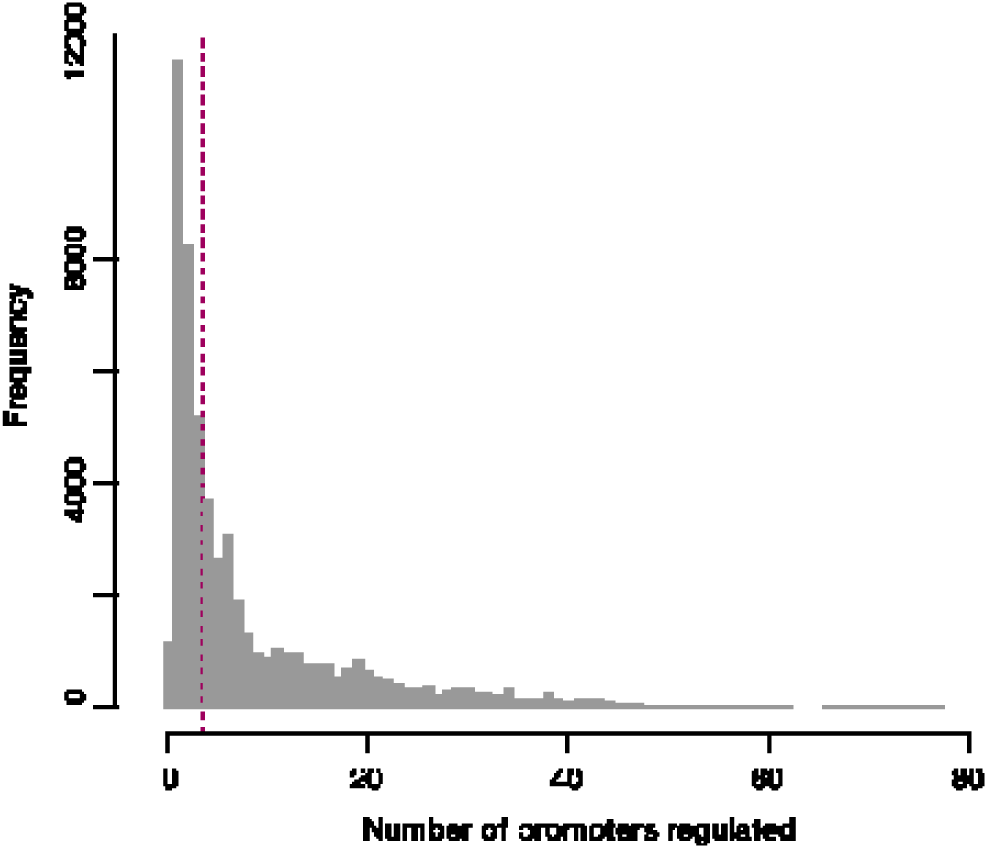
Number of genes in ECF σ regulons.

A key feature of global regulators is their importance under diverse conditions. To assess the importance of ECF σs with large regulons under various conditions, we used a previously published dataset (Price et al. 2018) in which 5,647 genome wide transposon fitness experiments were performed on 37 diverse bacterial species. A total of 136 ECF σs from our data set, 77 of which had regulon predictions, were represented in these experiments, allowing us to determine if predicted regulon size correlated to the number of conditions in which each ECF σ affects cellular fitness. We found that the number of regulated promoters was significantly (*p* < 0.0001) correlated to the fraction of conditions in which the ECF σ was important for cellular fitness (t < -3). On average, ECF σs predicted to regulate large regulons (>3 promoters) had significant phenotypes in 4.2% of tested conditions, whereas ECF σs with small regulons (≤3 promoters) had significant phenotypes in 0.4% of conditions (t-test *p* < 0.0001). Several RpoE and RpoE2 homologs in *Shewanella* species were predicted to regulate large regulons but exhibited few significant phenotypes. Since these ECF σs have been reported to be important under a wide range of growth and stress conditions (Dai et al. 2015), the lack of significant phenotypes is likely due to experimental issues (such as compensatory mutations). Notably, the regulons of these ECF σs appears to be correctly predicted. Another important characteristic of global regulators is their conservation within bacterial clades. To determine whether ECF σs with large (>3 promoters) predicted regulons are more conserved than ECFs with small (≤3 promoters) predicted regulons, we grouped ECF σs by their eggNOG identifiers and assessed in what fraction of genomes within a bacterial clade each eggNOG appeared. We found that ECF σs with large regulons were found on average in 28% of genomes within a clade, while ECF σs with small regulons were found on average in 16.9% of genomes within a clade. Due to the varying diversity within clades and within eggNOG groups, our measure of penetrance within clades likely underestimates the ultimate conservation of ECF σs. Despite this, the statistically significant (t-test *p <* 10^−5^) difference in distribution suggests that ECF σs with large regulons, such as RpoE in the γ-proteobacteria and SigR/SigH in the Actinobacteria, are conserved in a majority of species within those clades, while ECF σs with smaller regulons tend to have a patchier distribution.

One explanation for the patchier distribution of ECF σs with small regulons is that they were acquired through horizontal gene transfer (HGT) from other species. Because of the universally conserved nature of RNAP in bacteria, ECF σs may be ideal regulators for HGT genes. To determine whether ECF σs are commonly horizontally acquired, we applied tetranucleotide profiling (Becq, Churlaud, and Deschavanne 2010) to all genomes in our data-set to identify regions of atypical sequence content associated with HGT events and assessed whether ECF σs were present in these regions (i.e. horizontally acquired). We found that ECF σs were not more likely to be horizontally acquired than other genes in most bacterial phyla (Supplementary Table 8). However, in Bacteroidetes and Cytophagia (which are both members of the Fibrobacteres, Chlorobi, and Bacteroidetes super-phylum) ECF σs were more frequently horizontally acquired (21% and 18%, respectively) than the already relatively high HGT rate of all genes in these phyla (16% and 13%, respectively). Many ECF σs in Bacteroidetes have been implicated in the regulation of carbohydrate utilization pathways, raising the possibility that these ECF σs are being transferred as part of carbohydrate utilization loci. Across all clades, ECF σs with small (≤3 promoters) regulons were more likely to be horizontally acquired than ECF σs with large (>3 promoters) regulons (9.7% vs 7.9%, z-test *p* < 10^−12^), raising the possibility that these ECF σs are be transferred together with their regulon. Consistent with this idea, we found that many of the genes regulated by these ECF σs were located within 2kb of the ECF σ (25.0%, compared with 7.8% for ECF σs with >3 regulated promoters).

Taken together, these data suggest that ECF σs can play very different roles in bacterial gene regulatory networks. Some ECF σs appear to function as local regulators, controlling the expression of a few promoters often located nearby in the genome, likely in response to specific stimuli. These ECF σs contrast with other ECF σs which appear to function as global regulators, controlling the expression of many promoters and functioning to maintain cellular homeostasis under diverse conditions.

### The role of ECF σs in bacterial clades

In addition to providing a database of ECF σs regulon predictions across clades to guide future discovery, our analysis also revealed specific regulatory interactions of biological interest which confirm and extend previous analyses. In addition to the few such examples described here, the full data-set (available online) provides many more intriguing connections.

A major finding of our survey of ECF σ regulons was that almost half of the ECF σs function as local regulators. This regulatory function is exemplified by two broadly distributed groups of σs: ECF41 and ECF42 groups. Previous work had noted the conserved synteny of the genes surrounding these ECF σs, and had experimentally validated the small regulon of these ECF σs in a model organism (Wecke et al. 2012; Q. Liu, Pinto, and Mascher 2018). Our analyses support these findings and expand them to numerous bacterial clades, highlighting the local nature of these regulators. Our recapitulation of these results also highlights the high stringency of our ECF σ regulon prediction.

The Actinobacteria are an expansive, diverse phylum of high-GC Gram-positive organisms. Actinobacteria are canonically associated with soil ecosystems and many are also relevant in human health and disease: *Streptomyces* produce antibiotics, *Bifidobacteria* are microbiome constituents, and *Mycobacteria* are important pathogens. Despite their varied niches, all Actinobacteria contain a homologous ECF σ, known as SigR in *Streptomyces* species and SigH in *Mycobacterium* species, which responds to redox and translation stress. To better understand how evolution has shaped the regulon of this ECF σ to enable survival in these different niches, we examined which genes were putatively regulated by this ECF σ in the six major actinobacterial orders. SigH/SigR was predicted to regulate 22 to 61 genes, consistent with previous reports (Park, Lee, and Roe 2019; Sharp et al. 2016). We identified a core regulon consisting of 6 genes (*iscA, trxB, clpC, SCO3296, rbpA*, and *sigH/R*) which were significantly regulated in all 6 orders and an additional 18 genes predicted to be regulated in 4-5 orders. Many of these conserved regulon members encode classical oxidative stress response proteins, such as thioredoxins, members of the Clp complex, and Fe-S cluster repair proteins. The regulon of this ECF σ in the Corynebacteriales (which includes *Mycobacterium)* contains 6 unique genes. Intriguingly, 3 of these (Rv3054c, Rv3463c, and Rv1334) were among the most strongly activated genes in a recent over-expression study (Sharp et al. 2016). Both Rv3054c and Rv3463c (the two most SigH activated proteins in *M. tuberculosis*) encode reductases, and Rv1334 has been shown to be essential for survival in macrophages (Rengarajan, Bloom, and Rubin 2005). Although the functions of these genes remain to be determined, their integration into the SigH regulon of Corynebacteriales, but not of the other orders, suggests a role for these proteins in responding to redox and translation stresses unique to the Corynebacteriales, potentially signaling a role for these proteins in infection.

SigX, an important ECF σ in *Pseudomonas* species (Blanka et al. 2014), illustrates another aspect of regulon evolution. This ECFσ is important for growth, biofilm formation, and virulence. It is predicted to regulate 28 genes, many of which are involved in fatty acid synthesis and other membrane functions, consistent with previous work (Boechat et al. 2013; Blanka et al. 2014). SigX is also predicted to regulate nearby genes *cmpX*, a conserved transmembrane protein, and *oprF*, the major *Pseudomonas* outer membrane porin. Despite its importance in *Pseudomonas*, SigX is found only in a few other closely related γ-proteobacterial clades (*Paraglaciecola, Glaciecola*, and *Colwellia*). Surprisingly, a SigX homolog is found in *Flavobacterium*, a class in the Bacteroidetes phylum. In this clade, SigX is predicted to regulate nearby genes homologous to *cmpX* and *oprF* but not genes involved in fatty-acid metabolism. These regulatory patterns, combined with the atypical tetranucleotide profile of SigX in a majority (84%) of γ-proteobacterial genomes, suggest that SigX and neighboring genes may have been horizontally acquired by *Pseudomonas*, possibly from a member of the Flavobacteria. If this is the case, regulation of fatty acid synthesis by SigX in *Pseudomonas* may be due to a regulatory capture of those genes by SigX over a relatively short time span, raising important questions about the evolutionary pressures that led to this outcome.

### PERSPECTIVE

Predicting gene regulatory interactions from the amino-acid sequence of transcriptional regulators is a long-standing goal in biology. To do so requires both the ability to predict the sequence specificity of a novel regulator and the ability to determine significant interactions. Because of the difficulty of meeting both of these challenges, previous work has focused primarily on engineering modular regulatory proteins such as zinc-finger (ZFs) and transcription activator-like (TALs) rather than on *de novo* prediction of biological targets, a problem that remains largely unsolved. For example, despite decades of study on ZFs structure and specificity, the regulatory role of a majority of the 700+ ZFs in the human genome remain unknown (Dogan et al. 2020; Zuo et al. 2019). In this manuscript, we both solve the DNA-specificity code of ECF σs, a major class of bacterial regulators, and build a computational pipeline that enables us to use these rules to determine statistically significant regulons for ∼67% of bacterial ECF σs. The resulting dataset is the first of its kind: a comprehensive look at the role of a broadly distributed regulatory family.

To determine the rules that govern ECF σ specificity, we built a compendium of σs and their putative auto-regulatory promoters and used mutual information (MI) analysis to identify a few key conserved interacting residues. The conserved nature of specificity determining interactions is likely due to constraints on ECF σ evolution imposed by the need to maintain extensive interactions not only between σ and RNAP, but also, in many cases, between the σ and its negative regulator (anti-σ). These extensive interfaces likely limit the ability of σs to evolve novel DNA specificities through conformational changes, thereby forcing specificity diversification to occur at existing DNA-interacting residues. We validated this analysis and demonstrated its utility for rational engineering of ECF σ specificity by comprehensively mutating 3 amino acid positions and 4 promoter positions of interest and confirming that substitutions altered promoter specificity as predicted by our analysis. Manipulation of ECF σ specificity has exciting implications for tuning bacterial regulation to increase stress resistance or metabolic productivity in industrial settings (Tripathi, Zhang, and Lin 2014). To leverage our ability to predict ECF σ binding sites into an understanding of the biological function of ECF σs, we developed a phylogenetic foot-printing pipeline that leverages the conservation of functional DNA elements between closely related genomes to stringently identify functional ECF σ promoters. Predicted ECF σ regulated promoters were preceded by UP-elements, preferentially contained pyrimidine-purine TSS, and shared a 5’UTR length distribution with experimentally determined promoters in closely related model organisms. These analyses extended our knowledge of how ECF σ promoters are constructed. Previous work on *E. coli* RpoE (Rhodius, Mutalik, and Gross 2012) suggested that, in contrast to σ^70^ driven promoters (Estrem et al. 1998), RpoE promoters are more effectively activated by distal UP-element subsites than by proximal UP-element subsites. Consistent with this observation, we found that ECF σ promoters from all bacterial phyla contained predominantly distal sites (Figure 4C), suggesting that UP-element subsite location may be broadly important for partitioning the bacterial transcriptional space by minimizing basal σ^70^ transcription from ECF σ promoters. Structural data suggests that proximal subsites may be unable to efficiently activate transcription from ECF σ promoters due to the positioning of ECF σ4, which precludes direct stabilizing contacts with the αCTD (Lane and Darst 2006) that help stabilize σ^70^ (Chen, Tang, and Ebright 2003).

Although the regulons of most ECF σs were accurately predicted, FecI-like σs were over-represented among ECF σs without statistically significant predictions. Previous experimental studies have suggested that FecI-like ECF σs may function differently from other ECF σs. For example, FecI is unable to induce the transcription of its regulon without the presence of FecR (Braun, Mahren, and Ogierman 2003), and mutagenesis of the promoter recognized by *Pseudomonas aeruginosa* PvdS, a FecI-like ECF σ, suggests that the stringency of -35 element recognition is weak (Wilson and Lamont 2006). Taken together, these data and the lack of statistically significant predictions for FecI-like ECF σs suggests that their promoter recognition properties may substantially differ from those of canonical ECF σs and merit further study.

The ECF σ regulon predictions generated here are the first comprehensive study of the function of a regulator across the bacterial phylogeny. Our analysis of this data revealed the breadth of ECF σ regulation, the presence of conserved global regulators and variable local regulators, and opened a window into the evolution of ECF σ regulon membership in different niches. However, the approach demonstrated here, which combines co-variation analyses with powerful phylogenetic foot-printing tools, is not limited to ECF σs, but can be broadly applied to study protein-DNA interactions, bridging experimental and computational approaches. Together, these approaches will lead to mechanistic and conceptual understanding of bacterial gene regulatory network function and evolution.

## ACKNOWLEDGMENTS

We thank H. Li, M. Laub, P. Kiley, V. Rhodius, and members of the C.A.G. Lab for extensive helpful discussions.

## AUTHOR CONTRIBUTIONS

HT and CAG designed the study; HT and KSM built software; HT, EAC and KSM performed analyses; HT performed experiments; HT, HO, EAC, KSM, TJD and wrote and/or edited the manuscript.

## FIGURES

**Supplementary Figure 1.**
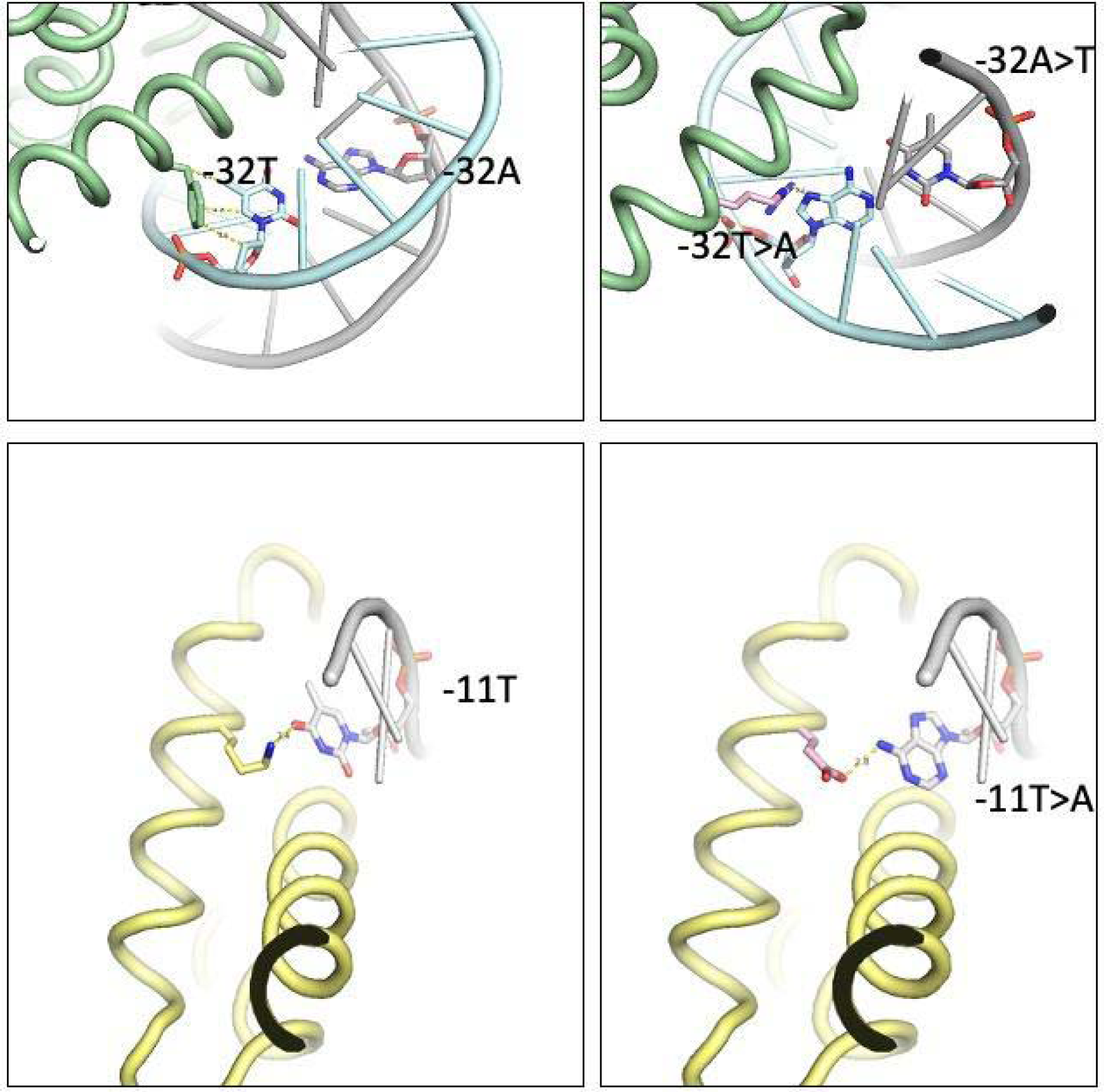
Structures of the Phe175-Arg175 and Lys56-Glu56 substitutions discussed in the text.

## METHODS

### Identifying ECF σs and their regulatory sequences

#### Identifying ECF σs

ECF σs from the ProGenomes database (Mende et al. 2017) were identified by using hmmer to search for protein sequences that contained both a σ2 domain (Pfam:Sigma70_r2, PF004542) and a σ4 domain (Pfam:Sigma70_r4, PF004545 or Pfam:Sigma70_r4_–_2, PF008281) but lacked a σ3 domain (Pfam:Sigma70_r3, PF004539), as previously described (Staroń et al. 2009). Sequences with more than 50 amino acids between the σ2 domain and the σ4 domain were excluded due to the likely presence of a cryptic σ3 domain. The resulting set contained 133,424 ECF σ sequences, which were aligned using Clustal Omega. Because alignments frequently contained gaps due to the sequence diversity of ECF σs, alignments were collapsed on either side of conserved residues in domains σ2.2, σ2.3, σ2.4, and σ4.2 to generate the final alignments (Supplementary Files 1,2).

#### Retrieving upstream sequences

To maximize the chance of finding conserved binding sites, up to 3 sequences at 2 lengths were retrieved for each ECF σ. For all ECF σs, sequences were retrieved upstream from translation start site of the ECF σ (1). If the ECF σ was in an operon (defined as co-directional ORFs < 50 nucleotides apart), the sequence upstream of the first gene in the operon was also retrieved (2). If a divergent gene was located either upstream of the ECF σ or of an operon containing the ECF σ, its upstream sequence was also retrieved (3). Sequences of both 150bp and 300bp were retrieved and analyzed, to increase the statistical power for small clusters (150bp) and to increase the probability of determining downstream motifs (300bp).

#### Duplicate removal

The genomes of certain organisms (e.g. *Mycobacterium tuberculosis, Streptococcus pneumoniae, Escherichia coli*) are grossly over-represented in sequence databases such as ProGenomes (Mende et al. 2017). To minimize the influence of duplicate sequences, we only considered one ECF σ from groups that were identical in protein sequence, upstream sequence, operon sequence, and divergent gene sequence. The remaining set contained 84,009 ECF σs.

#### Clustering

ECF σs were clustered solely on the basis of their protein sequence. We used two distinct methods to cluster ECF σs. First, we used the eggNOG identifier of each ECF σ. This clade specific identifier is based on all-by-all Smith-Waterman alignments of protein sequences from representative genomes (Huerta-Cepas et al. 2016). Our set contained 1,196 distinct eggNOG identifiers. Second, we used k-mer distance, as implemented in ClustalW. Besides the advantage of being a distinct clustering method, k-mer distance does not take into account phylogeny, allowing similar ECF σs from distinct bacterial clades to be interrogated together.

#### Motif discovery and weighting

For each ECF σ cluster, up to six libraries of upstream regulatory sequences were searched for putative 2-block motifs using BioProspector (X. Liu, Brutlag, and Liu 2001). These libraries consisted of 150bp and 300bp sequences from upstream of the ECF σs, upstream of an operon containing the ECF σs, and upstream of a divergently transcribed gene. Motif searches with BioProspector were performed only on the forward strand. Typical searches were of the form: W7 w5 G18 g15, where W and w denote the size (bp) of the -35 and -10 motifs, and G and g denotes the largest and smallest spacer size. For each library, the highest scoring 2-block motif was examined and aligned manually. Because an ECF σ could potentially be associated with as many as 12 motifs (150bp & 300bp; k-mer & eggNOG clusters; upstream, operon, & divergent sequences), or as few as 1, motifs were weighted by the reciprocal of the total number of motifs associated with each ECF σ (e.g. the only motif of an ECF σ has a weight of 1, whereas all 12 motifs of an ECF σ with 12 motifs are assigned weights of 1/12).

### Mutual information (MI) analysis

Mutual information analysis was performed on every possible nucleotide-position and amino-acid position pair.

### ECF σ substitution experiments

The σ-promoter GPF measurements were performed as previously described (Rhodius et al. 2013). Briefly, assays were performed in *E. coli* (DH10β) using a 3 plasmid system consisting of (1) an IPTG-inducible T7 RNAP (pN565), (2) a plasmid carrying the σ under the control of an T7 promoter, and (3) a plasmid series carrying a σ promoter driving to *sfgfp*. Assays were performed using 1mM IPTG, except those using ECF11_987, which were performed at 10μM IPTG to avoid toxicity. All assays were performed in a 96-well format. Overnight liquid transformants were diluted into fresh prewarmed LB +Spec, Amp, Kan and PTG in a 96-well cell-culture plate and covered with a breathable membrane. Cultures were incubated in a Tecan infinite M1000Pro plate shaker for 37°C at 582, r.p.m. OD600 and GFP fluorescence were tracked, and the maximum GFP synthesis rate/OD600 was calculated and used for downstream analysis.

### Modeling studies

Previously solved ECF structures (2MAP and 2H27) were modified using PyMOL and manually assessed for likely interactions.

### Phylogenetic foot-printing pipeline

A custom pipeline (github) was built to perform phylogenetic foot-printing based on eggNOG orthology (Huerta-Cepas et al. 2016). Input consisted of genomes of interest and the predicted - 35 and -10 PWMs for each genome. Briefly, each set of PWMs was scored on the upstream sequences (-332 to +2) of each gene in the appropriate genomes. Genes in operons were allowed to take the score of upstream genes, if higher. The score for each orthologous group (OG) was calculated by summing the score of the highest scoring member in each genome, weighted by the relative phylogenetic similarity of all genomes in the set. FDR thresholds were set by randomizing the order of the columns in the PWMs 200x and re-computing ortholog scores.

### Using gene expression patterns to evaluate regulon predictions

To evaluate the accuracy of the predicted regulons of ECF σs, regulons from 13 diverse bacterial species were used: *Mycobacterium tuberculosis H37Rv, Streptomyces coelicolor A3(2), Caulobacter crescentus CB15, Rhizobium etli CFN 42, Rhodobacter sphaeroides 2.4.1, Bacillus subtilize subsp subtilis str. 168, Bacteroides thetaiotaomicon VPI-5482, Porphyromonas gingivalis W83, Bordetella pertussis Tohama I, Burkholderia cenocepacia J2315, Escherichia coli str. K-12 substr W3110, Pseudomonas aeruginosa*, and *Shewanella oneidensis MR-1*. For each species, the program COnTORT (Myers et al. 2020) was used to download all publicly available gene expression data from NCBI GEO in order to capture as many unique experimental conditions as possible. Since the precise conditions required to activate each ECF σ is unknown, we used the large data set of all gene expression data with the idea that the more varied experimental conditions, the more likely the genes in the regulon would have correlated expression patterns. For each predicted regulon with at least two predicted members, the gene expression profile for all members of the regulon were correlated across all available gene expression data sets. To determine if the correlation was due to random chance, the analysis was performed using the same number of randomly selected genes (repeated 1,000 times). A Wilcoxon rank sum test in the software package R (version 3.6.1) was used to determine statistical significance between the correlation values of the genes within the predicted regulon and the randomly selected genes, with *p*-value of 0.05 being significant.

### Tetranucleotide profiling to identify HGT regions

Tetranucleotide profiling was performed as previously described (Becq, Churlaud, and Deschavanne 2010). Regions whose distance from the genomic median was > 2.5 MAD higher than normal were considered to have atypical sequence content (and therefore likely horizontally acquired).

## REFERENCES

Basu, Ritwika S., Brittany A. Warner, Vadim Molodtsov, Danil Pupov, Daria Esyunina, Carlos Fernández-Tornero, Andrey Kulbachinskiy, and Katsuhiko S. Murakami. 2014. “Structural Basis of Transcription Initiation by Bacterial RNA Polymerase Holoenzyme.” The Journal of Biological Chemistry 289 (35): 24549–59.

Beck, Heather J., and Isabella Moll. 2018. “Leaderless mRNAs in the Spotlight: Ancient but Not Outdated!” Microbiology Spectrum 6 (4). https://doi.org/10.1128/microbiolspec.RWR-0016-2017.

Becq, Jennifer, Cécile Churlaud, and Patrick Deschavanne. 2010. “A Benchmark of Parametric Methods for Horizontal Transfers Detection.” PloS One 5 (4): e9989.

Blanka, Andrea, Sebastian Schulz, Denitsa Eckweiler, Raimo Franke, Agata Bielecka, Tanja Nicolai, Fiordiligie Casilag, et al. 2014. “Identification of the Alternative Sigma Factor SigX Regulon and Its Implications for Pseudomonas Aeruginosa Pathogenicity.” Journal of Bacteriology 196 (2): 345–56.

Boechat, Ana Laura, Gilberto Hideo Kaihami, Mario José Politi, François Lépine, and Regina L. Baldini. 2013. “A Novel Role for an ECF Sigma Factor in Fatty Acid Biosynthesis and Membrane Fluidity in Pseudomonas Aeruginosa.” PloS One 8 (12): e84775.

Bonneau, Richard, Marc T. Facciotti, David J. Reiss, Amy K. Schmid, Min Pan, Amardeep Kaur, Vesteinn Thorsson, et al. 2007. “A Predictive Model for Transcriptional Control of Physiology in a Free Living Cell.” Cell 131 (7): 1354–65.

Braun, Volkmar, Susanne Mahren, and Monica Ogierman. 2003. “Regulation of the FecI-Type ECF Sigma Factor by Transmembrane Signalling.” Current Opinion in Microbiology 6 (2): 173–80.

Campagne, Sébastien, May E. Marsh, Guido Capitani, Julia A. Vorholt, and Frédéric H-T Allain. 2014. “Structural Basis for -10 Promoter Element Melting by Environmentally Induced Sigma Factors.” Nature Structural & Molecular Biology 21 (3): 269–76.

Chen, Hao, Hong Tang, and Richard H. Ebright. 2003. “Functional Interaction between RNA Polymerase α Subunit C-Terminal Domain and σ70 in UP-Element- and Activator-Dependent Transcription.” Molecular Cell 11 (6): 1621–33.

Dai, Jingcheng, Hehong Wei, Chunyuan Tian, Fredrick Heath Damron, Jizhong Zhou, and Dongru Qiu. 2015. “An Extracytoplasmic Function Sigma Factor-Dependent Periplasmic Glutathione Peroxidase Is Involved in Oxidative Stress Response of Shewanella Oneidensis.” BMC Microbiology 15 (February): 34.

Dogan, Berat, Senthilkumar Kailasam, Aldo Hernández Corchado, Naghmeh Nikpoor, and Hamed S. Najafabadi. 2020. “A Domain-Resolution Map of in Vivo DNA Binding Reveals the Regulatory Consequences of Somatic Mutations in Zinc Finger Transcription Factors.” bioRxiv, June. https://doi.org/10.1101/630756.

Donohue, Timothy J. 2019. “Shedding Light on a Group IV (ECF11) Alternative σ Factor.” Molecular Microbiology 112 (2): 374–84.

Estrem, S. T., T. Gaal, W. Ross, and R. L. Gourse. 1998. “Identification of an UP Element Consensus Sequence for Bacterial Promoters.” Proceedings of the National Academy of Sciences. https://doi.org/10.1073/pnas.95.17.9761.

Fang, Chengli, Lingting Li, Liqiang Shen, Jing Shi, Sheng Wang, Yu Feng, and Yu Zhang. 2019. “Structures and Mechanism of Transcription Initiation by Bacterial ECF Factors.” Nucleic Acids Research. https://doi.org/10.1093/nar/gkz470.

Fiebig, Aretha, Julien Herrou, Jonathan Willett, and Sean Crosson. 2015. “General Stress Signaling in the Alphaproteobacteria.” Annual Review of Genetics 49 (October): 603–25.

Gruber, Tanja M., and Carol A. Gross. 2003. “Multiple Sigma Subunits and the Partitioning of Bacterial Transcription Space.” Annual Review of Microbiology 57: 441–66.

Hubin, Elizabeth A., Mirjana Lilic, Seth A. Darst, and Elizabeth A. Campbell. 2017. “Structural Insights into the Mycobacteria Transcription Initiation Complex from Analysis of X-Ray Crystal Structures.” Nature Communications 8 (July): 16072.

Huerta, Araceli M., and Julio Collado-Vides. 2003. “Sigma70 Promoters in Escherichia Coli: Specific Transcription in Dense Regions of Overlapping Promoter-like Signals.” Journal of Molecular Biology 333 (2): 261–78.

Huerta-Cepas, Jaime, Damian Szklarczyk, Kristoffer Forslund, Helen Cook, Davide Heller, Mathias C. Walter, Thomas Rattei, et al. 2016. “eggNOG 4.5: A Hierarchical Orthology Framework with Improved Functional Annotations for Eukaryotic, Prokaryotic and Viral Sequences.” Nucleic Acids Research 44 (D1): D286–93.

Kelemen, Gabriella H., Gary L. Brown, Ján Kormanec, Laura Potúčkova, Keith F. Chater, and Mark J. Buttner. 1996. “The Positions of the Sigma-Factor Genes, whiG and sigF, in the Hierarchy Controlling the Development of Spore Chains in the Aerial Hyphae of Streptomyces Coelicolor A3 (2).” Molecular Microbiology 21 (3): 593–603.

Kim, Keon Young, Jeong Kuk Park, and Sangyoun Park. 2016. “In Streptomyces Coelicolor SigR, Methionine at the -35 Element Interacting Region 4 Confers the -31′-Adenine Base Selectivity.” Biochemical and Biophysical Research Communications 470 (2): 257–62.

Koo, Byoung-Mo, Virgil A. Rhodius, Gen Nonaka, Pieter L. deHaseth, and Carol A. Gross. 2009. “Reduced Capacity of Alternative Sigmas to Melt Promoters Ensures Stringent Promoter Recognition.” Genes & Development 23 (20): 2426–36.

Lane, William J., and Seth A. Darst. 2006. “The Structural Basis for Promoter-35 Element Recognition by the Group IV σ Factors.” PLoS Biology 4 (9): e269.

Li, Lingting, Chengli Fang, Ningning Zhuang, Tiantian Wang, and Yu Zhang. 2019. “Structural Basis for Transcription Initiation by Bacterial ECF σ Factors.” Nature Communications 10 (1): 1153.

Lin, Wei, Sukhendu Mandal, David Degen, Min Sung Cho, Yu Feng, Kalyan Das, and Richard H. Ebright. 2019. “Structural Basis of ECF-σ-Factor-Dependent Transcription Initiation.” Nature Communications 10 (1): 710.

Liu, Qiang, Daniela Pinto, and Thorsten Mascher. 2018. “Characterization of the Widely Distributed Novel ECF42 Group of Extracytoplasmic Function σ Factors in Streptomyces Venezuelae.” Journal of Bacteriology 200 (21). https://doi.org/10.1128/JB.00437-18.

Liu, X., D. L. Brutlag, and J. S. Liu. 2001. “BioProspector: Discovering Conserved DNA Motifs in Upstream Regulatory Regions of Co-Expressed Genes.” Pacific Symposium on Biocomputing. Pacific Symposium on Biocomputing, 127–38.

Luscombe, Nicholas M., M. Madan Babu, Haiyuan Yu, Michael Snyder, Sarah A. Teichmann, and Mark Gerstein. 2004. “Genomic Analysis of Regulatory Network Dynamics Reveals Large Topological Changes.” Nature 431 (7006): 308–12.

Martínez-Antonio, Agustino, and Julio Collado-Vides. 2003. “Identifying Global Regulators in Transcriptional Regulatory Networks in Bacteria.” Current Opinion in Microbiology 6 (5): 482–89.

Mecsas, J., P. E. Rouviere, J. W. Erickson, T. J. Donohue, and C. A. Gross. 1993. “The Activity of Sigma E, an Escherichia Coli Heat-Inducible Sigma-Factor, Is Modulated by Expression of Outer Membrane Proteins.” Genes & Development 7 (12B): 2618–28.

Mende, Daniel R., Ivica Letunic, Jaime Huerta-Cepas, Simone S. Li, Kristoffer Forslund, Shinichi Sunagawa, and Peer Bork. 2017. “proGenomes: A Resource for Consistent Functional and Taxonomic Annotations of Prokaryotic Genomes.” Nucleic Acids Research 45 (D1): D529–34.

Mendoza-Vargas, Alfredo, Leticia Olvera, Maricela Olvera, Ricardo Grande, Leticia Vega-Alvarado, Blanca Taboada, Verónica Jimenez-Jacinto, et al. 2009. “Genome-Wide Identification of Transcription Start Sites, Promoters and Transcription Factor Binding Sites in E. Coli.” PloS One 4 (10): e7526.

Myers, Kevin S., Michael Place, Daniel R. Noguera, and Timothy J. Donohue. 2020. “COnTORT: COmprehensive Transcriptomic ORganizational Tool for Simultaneously Retrieving and Organizing Numerous Gene Expression Data Sets from the NCBI Gene Expression Omnibus Database.” Microbiology Resource Announcements 9 (25). https://doi.org/10.1128/MRA.00587-20.

Paget, Mark S. 2015. “Bacterial Sigma Factors and Anti-Sigma Factors: Structure, Function and Distribution.” Biomolecules 5 (3): 1245–65.

Park, Joo-Hong, Ju-Hyung Lee, and Jung-Hye Roe. 2019. “SigR, a Hub of Multilayered Regulation of Redox and Antibiotic Stress Responses.” Molecular Microbiology 112 (2): 420–31.

Price, Morgan N., Kelly M. Wetmore, R. Jordan Waters, Mark Callaghan, Jayashree Ray, Hualan Liu, Jennifer V. Kuehl, et al. 2018. “Mutant Phenotypes for Thousands of Bacterial Genes of Unknown Function.” Nature. https://doi.org/10.1038/s41586-018-0124-0.

Rengarajan, Jyothi, Barry R. Bloom, and Eric J. Rubin. 2005. “Genome-Wide Requirements for Mycobacterium Tuberculosis Adaptation and Survival in Macrophages.” Proceedings of the National Academy of Sciences of the United States of America 102 (23): 8327–32.

Rhodius, Virgil A., and Vivek K. Mutalik. 2010. “Predicting Strength and Function for Promoters of the Escherichia Coli Alternative Sigma Factor, σE.” Proceedings of the National Academy of Sciences of the United States of America 107 (7): 2854–59.

Rhodius, Virgil A., Vivek K. Mutalik, and Carol A. Gross. 2012. “Predicting the Strength of UP-Elements and Full-Length E. Coli σE Promoters.” Nucleic Acids Research 40 (7): 2907–24.

Rhodius, Virgil A., Thomas H. Segall-Shapiro, Brian D. Sharon, Amar Ghodasara, Ekaterina Orlova, Hannah Tabakh, David H. Burkhardt, et al. 2013. “Design of Orthogonal Genetic Switches Based on a Crosstalk Map of σs, Anti-σs, and Promoters.” Molecular Systems Biology 9 (October): 702.

Rhodius, Virgil A., Won Chul Suh, Gen Nonaka, Joyce West, and Carol A. Gross. 2006. “Conserved and Variable Functions of the sigmaE Stress Response in Related Genomes.” PLoS Biology 4 (1): e2.

Sharp, Jared D., Atul K. Singh, Sang Tae Park, Anna Lyubetskaya, Matthew W. Peterson, Antonio L. C. Gomes, Lakshmi-Prasad Potluri, Sahadevan Raman, James E. Galagan, and Robert N. Husson. 2016. “Comprehensive Definition of the SigH Regulon of Mycobacterium Tuberculosis Reveals Transcriptional Control of Diverse Stress Responses.” PloS One 11 (3): e0152145.

Staron, Anna, Heidi J. Sofia, Sascha Dietrich, Luke E. Ulrich, Heiko Liesegang, and Thorsten Mascher. 2009. “The Third Pillar of Bacterial Signal Transduction: Classification of the Extracytoplasmic Function (ECF) σ Factor Protein Family.” Molecular Microbiology 74 (3): 557–81.

Tripathi, Lakshmi, Yan Zhang, and Zhanglin Lin. 2014. “Bacterial Sigma Factors as Targets for Engineered or Synthetic Transcriptional Control.” Frontiers in Bioengineering and Biotechnology 2 (September): 33.

Urtecho, G., K. Insigne, A. D. Tripp, M. Brinck, and N. B. Lubock. 2020. “G Enome-Wide Functional Characterization Of Escherichia Coli P Romoters and Regulatory Elements Responsible for Their Function.” bioRxiv. https://www.biorxiv.org/content/10.1101/2020.01.04.894907v1.abstract.

Wecke, Tina, Petra Halang, Anna Staroń, Yann S. Dufour, Timothy J. Donohue, and Thorsten Mascher. 2012. “Extracytoplasmic Function σ Factors of the Widely Distributed Group ECF41 Contain a Fused Regulatory Domain.” MicrobiologyOpen 1 (2): 194–213.

Wilson, Megan J., and Iain L. Lamont. 2006. “Mutational Analysis of an Extracytoplasmic-Function Sigma Factor to Investigate Its Interactions with RNA Polymerase and DNA.” Journal of Bacteriology 188 (5): 1935–42.

Zuo, Zheng, Timothy Billings, Michael Walker, Petko Petkov, Polly Fordyce, and Gary D. Stormo. 2019. “Why Do Long Zinc Finger Proteins Have Short Motifs?” bioRxiv. https://doi.org/10.1101/637298.

